# PhytoNet: Comparative co-expression network analyses across phytoplankton and land plants

**DOI:** 10.1101/255067

**Authors:** Camilla Ferrari, Sebastian Proost, Colin Ruprecht, Marek Mutwil

## Abstract

Phytoplankton consists of autotrophic, photosynthesizing microorganisms that are a crucial component of freshwater and ocean ecosystems. However, despite being the major primary producers of organic compounds, accounting for half of the photosynthetic activity worldwide and serving as the entry point to the food chain, functions of most of the genes of the model phytoplankton organisms remain unknown. To remedy this, we have gathered publicly available expression data for one chlorophyte, one rhodophyte, one haptophyte, two heterokonts and four cyanobacteria and integrated it into our PlaNet (Plant Networks) database, which now allows mining gene expression profiles and identification of co-expressed genes of 19 species. We exemplify how the co-expressed gene networks can be used to reveal functionally related genes and how the comparative features of PhytoNet allow detection of conserved transcriptional programs between cyanobacteria, green algae, and land plants. Additionally, we illustrate how the database allows detection of duplicated transcriptional programs within an organism, as exemplified by two DNA repair programs within *Chlamydomonas reinhardtii*. PhytoNet is available from www.gene2function.de.

## Introduction

While phytoplankton (used here in the broadest sense that includes cyanobacteria) is often not visible by unaided eye, it plays a crucial role in marine environments. These photosynthesis-capable organisms form the foundation of aquatic ecosystems and contribute to roughly 50% of the global carbon assimilation (1–3). Cyanobacteria are extant relatives of the organisms that gave rise to plant chloroplasts through endosymbiosis ~1.5 billion years ago, and therefore are vital to understand the origin of plants (4). Furthermore, the phylogenetic position of green algae makes them interesting outgroups for evolutionary studies focussed on higher plants (5). Having often a simpler genetic makeup than higher plants, various single-celled algae have been hailed as a “green yeasts”, with potential applications ranging from the production of biofuels to food supplements (6, 7). Finally, diatoms and coccolithophores, which construct intricate shells from silica and calcium carbonate respectively (8, 9), have value for biotechnology and nanotechnology (10).

Despite their importance, the function of many algal genes remains unknown, and gene function prediction approaches for algae rely mainly on sequence similarity analysis. These approaches thus lag behind when compared to gene function prediction methods applied to model organisms in other branches of the tree-of-life (11). Online resources focussed on algae are rare, and while a comparative genomics platform was established recently (12), tools to study gene expression in algae are still lacking. As genes with similar expression profiles across developmental stages, environmental perturbations and organs tend to be functionally related, identification of co-expressed genes has been used to reveal functionally related genes (13–17). These co-expression relationships can be visualized as networks, where nodes (or vertices) correspond to genes and edges (or links) connect genes that display similar expression profiles (18). Thus, by identifying groups of connected genes in these networks, functional gene modules can be detected and used to predict gene function. Furthermore, gene modules conserved over large phylogenetic distances can reveal core components of shared processes, while duplicated gene modules can reflect recent adaptations combined with an increase in complexity (19–21).

Here we present PhytoNet (www.gene2function.de), a freely available extension to PlaNet, which allows studying co-expression networks in nine algae through an online interface.

## Materials and Methods

### Integration of nine phytoplankton species into the PlaNet database

To make PhytoNet a complete platform to study gene expression of phytoplankton, we screened ArrayExpress database for expression data of photosynthesizing microorganisms (22), with criteria: (i) the candidate organism must be sequenced and (ii) at least 20 expression datasets must be available (23, 24). We arrived at nine species (Table 1, Figure 1A, Supplementary Table 1), which include four cyanobacteria (*Synechocystis sp. PCC 6803*, *Nostoc punctiforme PCC 73102*, *Prochlorococcus marinus subsp. pastoris str. CCMP1986*, *Cyanothece sp. ATCC 51142*), two heterokonts (*Ectocarpus siliculosus*, *Phaeodactylum tricornutum*), one rhodophyte (*Cyanidioschyzon merolae*), one haptophyte (*Emiliania huxleyi*) and one chlorophyte (*Chlamydomonas reinhardtii*). Synechocystis, Cyanothece, Phaeodacticum and Emiliania raw microarray data was downloaded and processed using the R-package limma (25), Nostoc and Ectocarpus raw data was normalized using Deva software (Roche, Nimblegen), Chlamydomonas raw RNA sequencing data was downloaded and transcript per million (TPM)-normalized using LSTrAP software (26), while Prochlorococcus and Phaeodacticum processed data was downloaded from ArrayExpress (Table S1). LSTrAP was used to detect and discard samples that showed either (i) low mapping to the genome (< 65%), (ii) low mapping to coding sequences (< 40%) or (iii) with too few useful reads (less than 8M reads mapping to the genome). The remaining samples were used to construct expression matrices and co-expression networks. Pfam domain labels were assigned to genes by blasting CDS sequences against Pfam database version 27 with an e-value cutoff of 10^-5^ (21, 27), while Gene Ontology terms and gene families were obtained by using the CDS sequences in TRAPID (28).

**Figure 1.**
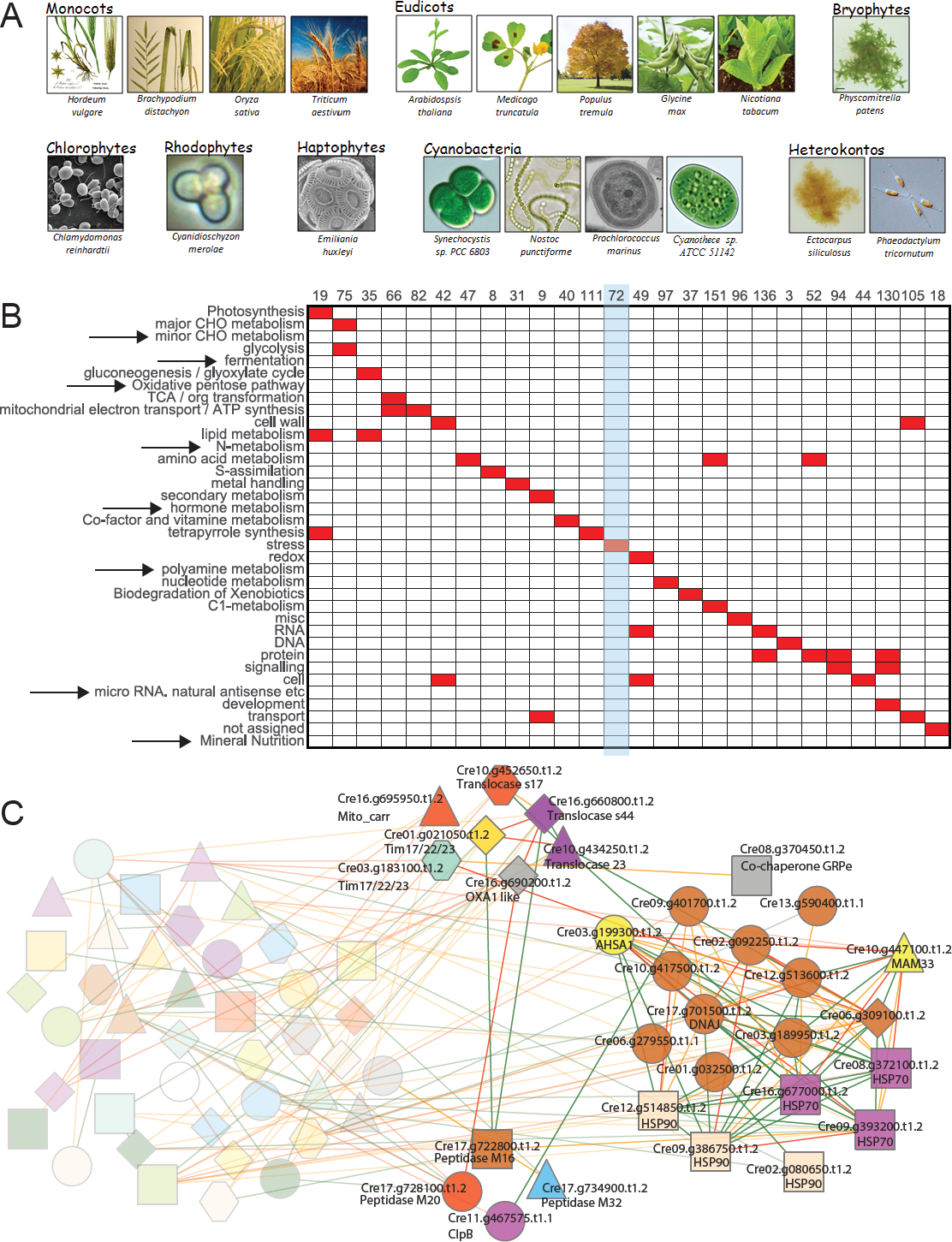
MapMan cluster analysis of *Chlamydomonas reinhardtii*. A) An overview of species found in PhytoNet. B) MapMan analysis of Chlamydomonas clusters. Rows represent MapMan bins (i.e., functional categories) while columns represent clusters. Red squares indicate that the specific term is significantly enriched (FDR-adjusted p-value < 0.05) in a given cluster. Arrows indicate bins with no significant enrichment. The light blue rectangle indicates the stress cluster 72. C) Co-expression network representation of cluster 72 (stress). Nodes represent genes while edges connect genes which are coexpressed. Colored shapes are used to indicate genes that belong to the same gene family and/or contain same Pfam domains. Colored edges indicate the degree of co-expression between genes with green, orange and red edges denoting Highest Reciprocal Rank (HRR) of 10, 20 and 30, respectively. For clarity, only discussed genes are highlighted.

**Table 1.**
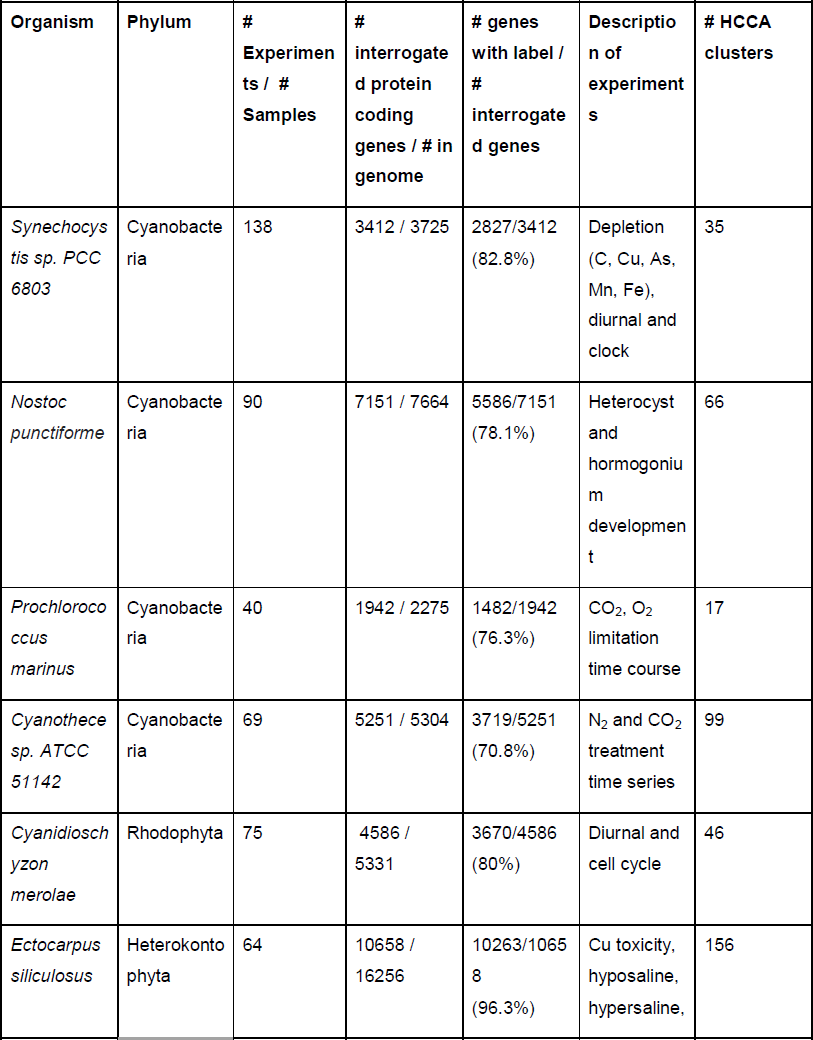
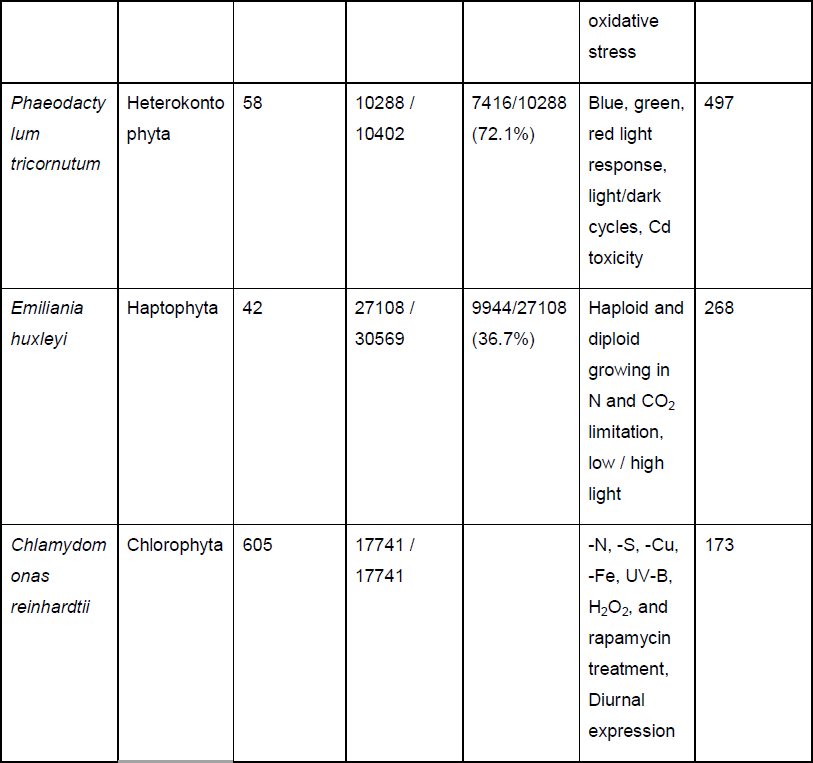
Description of the utilized expression data. The table indicates the number of expression measurements (samples), fraction of total protein-coding genes captured by the expression measurement platform (microarray or RNA sequencing), fraction of interrogated protein-coding genes with assigned Pfam domain and/or PLAZA gene family, types of experiments and the number of HCCA clusters.

To assess the quality of the expression data, we first analyzed the sample similarity dendrogram, with the assumption that replicates and samples representing similar experiments should group together. The sample similarity dendrograms showed an expected clustering of replicates and related samples (i.e., separate clustering of Synechocystis day and night samples, Supplementary Figure 1-9). Furthermore, co-expression networks tend to show a scale-free topology (also called power law behavior), where most nodes have few connections, and few nodes have many connections (29). The expression data of the nine species produced the expected pattern on a line with a negative slope (recommended Pearson Correlation Coefficient cutoff >0.7, (30), Supplementary Figure 1-9), which is indicative of scale-freeness of the co-expression networks (24, 31, 32). Taken together, this indicates that the data is of sufficient quantity and quality to produce co-expression networks with correct topology.

The expression data was used to construct Highest Reciprocal Rank (HRR) co-expression networks, and the clusters of co-expressed genes were identified with Heuristic Cluster Chiseling Algorithm (HCCA, (32)). To detect conserved and duplicated modules, we applied the FamNet pipeline, which identifies co-expression neighborhoods that contain same gene families and Pfam domains (21). Briefly, when two neighborhoods of sizes A and B are compared, the number of Pfam domains and gene families (labels) in common between the two neighborhoods is counted and compared to an expected number of random neighborhoods of these sizes (21). Finally, the data was uploaded to the PlaNet database (33).

## Results

### Identification of functionally related gene clusters - protein folding

In order to have a clear overview of the biological processes present in the co-expression networks, the networks were clustered with the HCCA algorithm, which is optimized to cluster HRR-based co-expression networks (32). For the unicellular green algae *Chlamydomonas reinhardtii*, 173 clusters were identified (the number of clusters identified for each species is shown in Table 1). In order to elucidate the biological functions of these clusters, we computed enriched MapMan gene ontology bins present in the clusters (34). 63 of the clusters showed significant enrichment in at least one MapMan bin, and 14 of them showed enrichment in more than one process (false discovery rate-adjusted p-value < 0.05, Figure 1B). Processes such as minor carbohydrate metabolism, fermentation, oxidative pentose pathway, nitrogen metabolism, hormone metabolism, polyamine metabolism, micro RNA and mineral nutrition were not enriched in any of the Chlamydomonas clusters, while the bin “protein” was present multiple times, indicating the presence of numerous bins involved in synthesis, modification, and degradation of proteins.

We demonstrate how these clusters can be used to identify relevant genes with a stress-related cluster from Chlamydomonas. Abiotic stress can negatively affect biomass production in all living organisms, and the algal response to stresses is being studied with the aim to transfer the obtained knowledge to food crops. Numerous studies have investigated the transcription profiles and physiological changes in Chlamydomonas caused by abiotic stresses (35–37). Consequently, we decided to explore cluster 72 (http://aranet.mpimp-golm.mpg.de/responder.py?name=gene!cre!c72), which showed significant enrichment for the MapMan bin “stress” (Figure 1B). Within the cluster, we found three groups of genes involved in three different processes (Figure 1C). The first and largest group comprised chaperones belonging to Pfams DNAJ (brown circle), HSP90 (beige square), HSP70 (violet square), chaperone activators AHSA1 (yellow circle, activating HSP90)(38) and GRPe (gray square, activating HSP70)(39). The second group contained metallopeptidases M16 (brown square), M20 (red circle), M32 (blue triangle) and a Clp ATPase involved in protein unfolding (ClpB, purple circle)(40). The third group included proteins involved in mitochondrial protein import, such as Translocases s17 (red hexagon), 23 (purple triangle), s44 (purple diamond), inner membrane translocases belonging to TIM17/22/23 families (yellow diamond, turquois hexagon), mitochondrial carriers translocating proteins across membranes (red triangle)(41) and OXA1, which is involved in protein insertion into the inner mitochondrial membrane (grey diamond)(42). These results revealed known components of protein folding, degradation and import into mitochondria, and suggest that these processes might be coordinated in Chlamydomonas. Taken together, the functionally enriched clusters can serve as a rapid entry point to identify genes involved in the biological process of interest.

### Detecting conserved gene modules - primary metabolism

The combination of gene families, Pfam domains, and co-expression modules allows detection and analysis of gene modules that are present in multiple species (13, 43, 44), and this property can be used to transfer functional knowledge from one species to another (45–48). Such conserved modules indicate that the biological process captured is present in the investigated species, and provides additional confidence that the homologous genes present in these modules are needed for the process and perform the same function (45).

To illustrate how conserved modules can be detected and analyzed, we used Synechocystis threonine synthase gene *sll1172* (http://aranet.mpimp-golm.mpg.de/responder.py?name=gene!syn!3362). The expression profile of *sll1172* showed that the gene is expressed in all conditions, and co-expressed with other genes involved in amino acid biosynthesis, glycolysis and nucleotide metabolism (the co-expression network and gene ontology enrichment analysis is found on the gene page). The database visualizes conserved modules by using a gene module network (Figure 2A). The nodes in the network represent modules (groups of co-expressed genes), blue edges indicate conserved modules (i.e. modules that contain same gene families and Pfam domains), while the edge style indicates the number of gene families and Pfam domains in common between two modules (49). Finally, modules that are overlapping (i.e. contain same co-expressed genes) are connected by orange edges, while green edges indicate duplicated modules (discussed in the next section).

The gene module network for *sll1172* indicated that conserved modules were found in flowering plants (green and orange nodes), cyanobacteria (blue nodes), early land plants (moss Physcomitrella, yellow nodes), red alga (Cyanidioschyzon, red nodes), green algae (Chlamydomonas, brown nodes), heterokonts (purple nodes) and haptophytes (Emiliania, gray nodes), suggesting that the module is conserved over large evolutionary distance. To exemplify a typical analysis and to gain insight into the function of these conserved modules, we selected Synechocystis *sll1172*, Arabidopsis *At3g59760* and Physcomitrella *Pp1s370_61v6.1* from the table below the module network and clicked on “Compare” button. The output of the analysis indicated which common gene families and Pfam domains were found in the three modules, by using colored shapes to show presence or absence of a family (Figure 2B). The database also provides a more detailed view of the modules, where individual genes are shown (Figure 2C). Closer inspection of the Arabidopsis module revealed numerous genes involved in ATP synthesis (indicated by a black box), amino acid synthesis (blue box), steroid biosynthesis (red box), glycolysis (green box) and adenylate salvage (purple box, Figure 2C legend). To conclude, these results show that the observed primary metabolic processes are transcriptionally coordinated in cyanobacteria, plantae and species that arose via secondary endosymbiosis, indicating that the observed modules are ancient.

**Figure 2.**
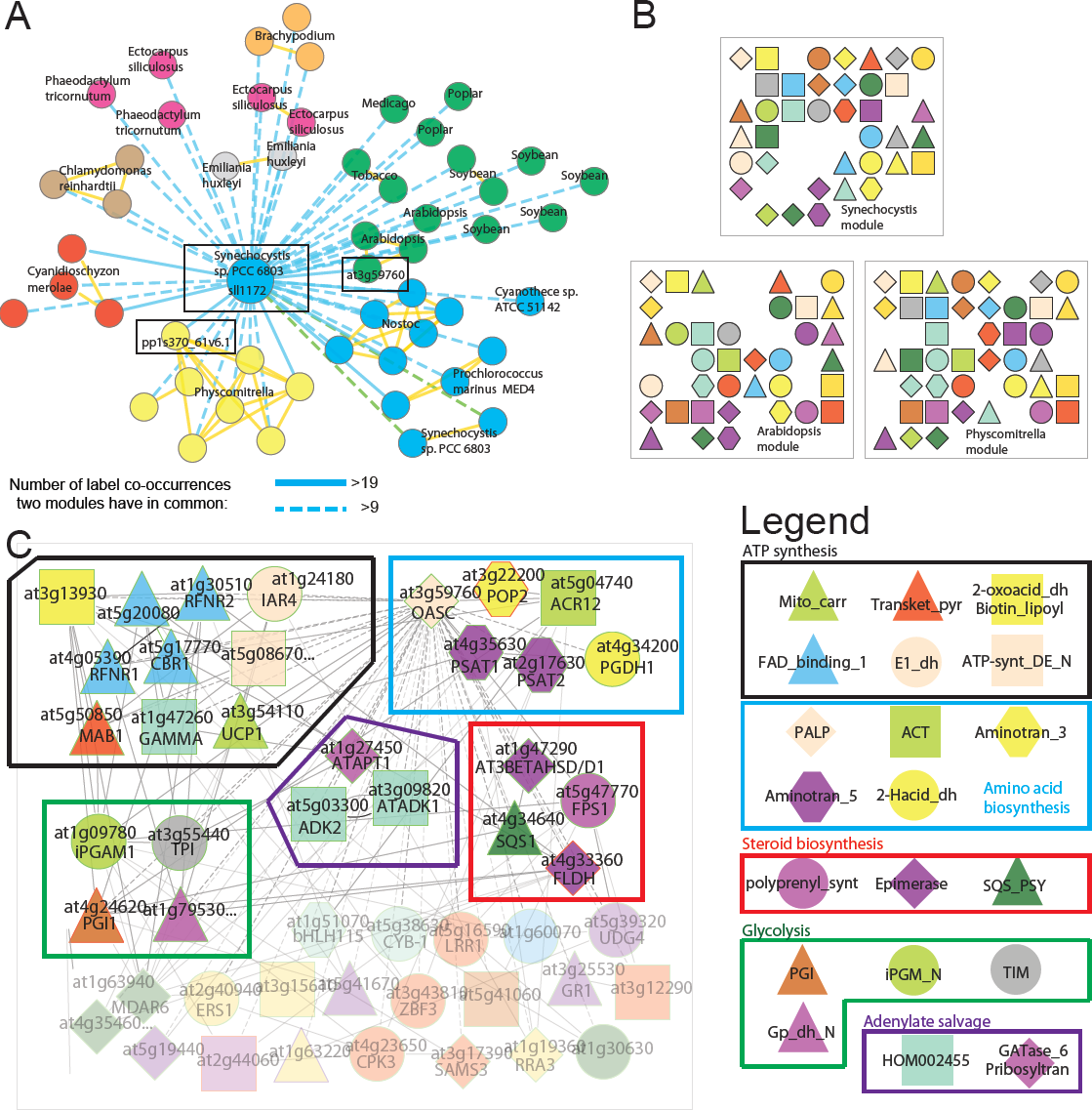
Detailed gene module analysis of the algae. A) Gene module network of *sll1172* showing only modules with at least 10 gene families and Pfams domains (= label co-occurrences) in common with the bait module. The central node (large, blue circle) represents *sll1172* gene module, while similar modules are connected by blue edges, where edges with heavier weight (solid edges) connect gene modules with at least 20 label co-occurrences in common. Yellow and green edges indicate overlapping modules and similar modules within the same species. The different phyla present in the network are indicated by the different node colors. For example, cyanobacteria, dicots, and bryophytes are indicated by blue, green and yellow nodes. The three selected modules are indicated by black boxes. B) Gene families and Pfam domains indicated by colored shapes, that the three modules have in common. C) Co-expression network of the Arabidopsis gene module. Nodes represent genes, colored shapes represent gene families and Pfam domains, while co-expressed genes are connected by gray edges. The discussed genes and gene families are highlighted by colored boxes.

### Duplicated gene modules - DNA repair

We have previously shown that duplication of co-expression modules is a widespread phenomenon in flowering plants and in early land plants, such as mosses (20, 45, 50). We exemplify such duplicated transcriptional modules in algae with *Chlamydomonas reinhardtii* gene *Cre01.g048200.t1.1* as a bait gene (http://aranet.mpimp-golm.mpg.de/responder.py?name=gene!cre!10456). This gene is annotated as a putative RNA helicase and contains “AAA” Pfam domains, which correspond to “ATPases associated with diverse cellular activities”. The gene module network for this gene revealed six highly similar modules that were overlapping with each other (Fig. 3A). Of these six modules, we selected *Cre03.g187950* for comparison with *Cre01.g048200*, as this showed the highest number of gene families and Pfam domains (expressed as label co-occurrences)(21) in common with our bait gene.

**Figure 3.**
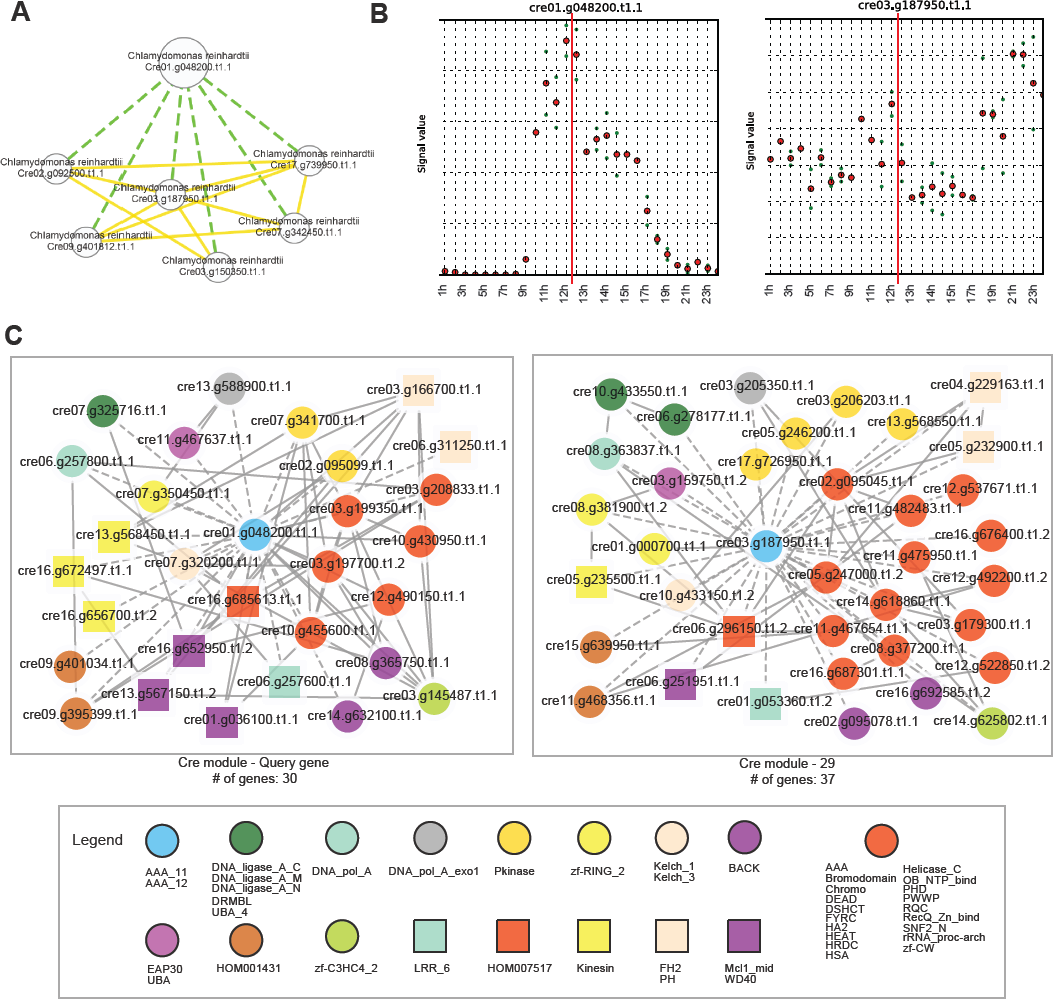
Duplicated modules involved in DNA repair in *Chlamydomonas reinhardtii*. (**A**) Gene module network of *Cre01.g048200* showing only Chlamydomonas modules sharing at least 10 label co-occurrences. Green edges indicate similarity of modules within the same species, whereas yellow edges indicate overlapping modules (**B**) Expression profiles of *Cre01.g048200* and *Cre03.g187950* during diurnal cycle (12h light/12h dark). The red line indicates the shift from light to dark. Full expression profiles can be viewed on the respective gene pages. (**C**) Network comparer result showing the similarity of *Cre01.g048200* and *Cre03.g187950* networks, with the query genes nodes enlarged. The legend provides label names and their corresponding functions in brackets. The corresponding full modules can be viewed at the respective gene pages at PhytoNet.

The expression patterns of these two genes are distinct, as *Cre03.g187950* is expressed during the whole diurnal cycle, whereas *Cre01.g048200* is induced at the end of the day with again decreasing expression during the night (Figure 3B). The network comparison of these two genes revealed common Pfam domains, such as “DNA_ligase_A”, “DNA_pol_A”, “Helicase_C”, and “DNA_pol_A_exo” (Figure 3C), which are associated to DNA repair, replication and proofreading, as well as several DNA/chromatin-associated Pfam domains, such as “Bromodomain”, and “SNF_2”. To discover the putative function of the two modules, we further analyzed the GO term enrichment of their respective individual gene networks. Interestingly, we found significant enrichment for genes involved in DNA repair in both modules and for DNA replication in the module that is induced at the end of the day (Supplementary Tables 2 and 3). In synchronized cell cultures under a 12 hour light/dark regime, *Chlamydomonas reinhardtii* cells divide with the onset of the night (51). As the DNA replication occurs shortly before, the module that is induced at the end of the day might be involved in DNA mismatch repair during DNA replication. Conversely, the constitutively expressed module might function as a general DNA repair module. Taken together, our analyses identified putatively duplicated DNA repair modules with distinct expression patterns and specialized functions.

## Conclusion

To remedy the lack of transcriptome analysis tools for phytoplankton, we introduce PhytoNet, an extension of the PlaNet database which adds cyanobacteria and eukaryotic algae to the web server, thus increasing the number of species to 19. The web server contains gene expression profile plots, gene co-expression networks and Gene Ontology enrichment analyses for gene neighborhoods and clusters, which can be used to predict gene functions. Furthermore, the comparative features of PlaNet allow detection of conserved and duplicated gene modules, allowing rapid identification of functionally related gene modules across and within phytoplankton and land plants. Therefore, PhytoNet is a versatile and easy to use hypothesis generation server for phytoplankton researchers who study gene functions in these organisms.

## Data availability

The expression matrices, co-expression networks, functionally enriched clusters and conserved/duplicated modules are found at http://aranet.mpimp-golm.mpg.de/download.html.

## Supplementary data

Supplementary Data are available at NAR online.

## Confict of interest

None.

